# High-NA open-top selective-plane illumination microscopy for biological imaging

**DOI:** 10.1101/140418

**Authors:** Ryan McGorty, Dan Xie, Bo Huang

## Abstract

Selective-plane illumination microscopy (SPIM) provides unparalleled advantages for volumetric imaging of living organisms over extended times. However, the spatial configuration of a SPIM system often limits its compatibility with many widely used biological sample holders such as multi-well chambers and plates. To solve this problem, we developed a high numerical aperture (NA) open-top configuration that places both the excitation and detection objectives on the opposite of the sample coverglass. We carried out a theoretical calculation to analyze the structure of the system-induced aberrations. We then experimentally compensated the system aberrations using adaptive optics combined with static optical components, demonstrating near-diffraction-limited performance in imaging fluorescently labeled cells.

© 2017 Optical Society of America

**OCIS codes:** (080.080) Geometric Optics; (110.0110) Imaging systems; (110.0180) Microscopy.

## 1. Introduction

For over a decade selective-plane illumination microscopy (SPIM), or light-sheet microscopy, has been successful in imaging the development of whole organisms [1, 2], tracking cell lineages in living embryos [3], observing individual neurons fire across large volumes [4], and observing the dynamics of single cells [5, 6]. SPIM systems often require special sample loading approaches, in which small tubes or cylinders of agarose gel hold the sample in the typically tight space between two objectives. This configuration makes it difficult for SPIM systems to use conventional sample holders for cell microscopy, particularly for high-content imaging where microfluidic devices or multi-well plates would be the ideal. Efforts have been devoted to resolve this incompatibility between SPIM systems and high-throughput, high-resolution imaging, however, the published methods are either optically demanding [7] or require a specially manufactured sample holder wherein the specimen is supported by a thin membrane [8]. A more user-friendly design which resembles typical inverted microscopes and suits existing popular sample holders has yet to be developed.

Previously, we demonstrated an open-top SPIM configuration to work with a variety of sample holders [9]. The two objectives, one for creating the excitation light sheet and the other for imaging, are installed underneath the sample holder (Fig. 1(a)). Mounting of the imaging sample in the open-top SPIM is similar to that in an inverted microscope, where the sample is carried by a flat glass surface such as a coverslip or the glass bottom of a multi-well plate. However, this convenience of sample loading comes at the expense of image quality. Because a piece of glass tilted with respect to the optical axis is between the objective and the sample, severe optical aberrations reduce the image quality. If the open-top SPIM uses a low numerical aperture (NA) imaging objective (e.g., NA = 0.3), the geometry-based aberration would primarily consist of spherical aberration and astigmatism, both of which are analytically simple. We employed two elements to help counter these aberrations: the first was a water prism to replace the tilted air-glass-water interface with a tilted water-glass-water interface. The second measure was a cylindrical lens placed in the imaging path. With these correction methods, a resolution of about 1 *μ*m was achieved using a 0.3 NA objective in our open-top SPIM. While micron-level SPIM resolution is adequate in many applications, it falls short of revealing details at the subcellular level. Therefore, it is desirable to use a higher NA objective for better resolution in our open-top SPIM system. However, such a goal cannot be achieved by simply replacing the imaging objective: under the published open-top SPIM setup, a high-NA objective would exacerbate the optical aberrations, which would go beyond the correction capacity of the water prism and cylindrical lens.

**Fig. 1.**
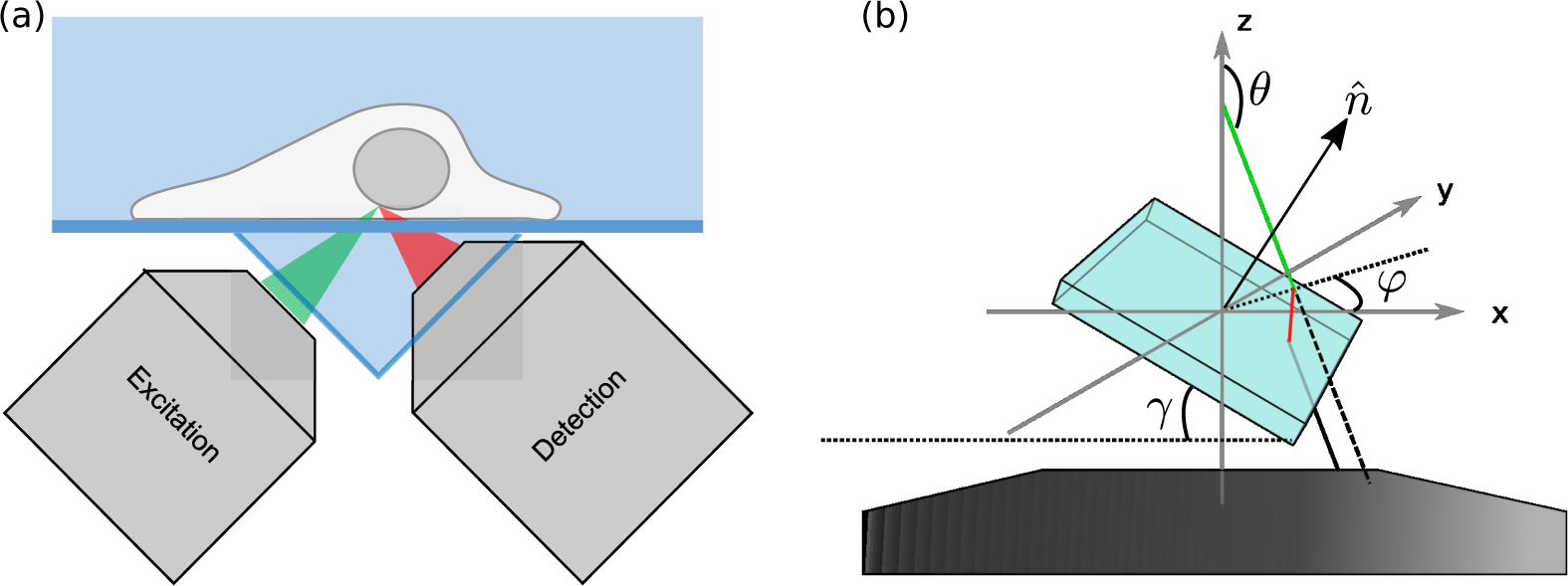
(a) The layout of the excitation and imaging paths of our previously described open-top SPIM. The sample sits on top of a cover-glass bottomed holder which is carried by a motorized stage. To allow for the water-dipping imaging objective and to partially mitigate the effects of imaging across a tilted coverslip, a water-holding triangular prism is placed directly underneath the sample holder. Note that the optical layout used for collecting the images and data shown here is modified to more easily understand and to test methods for compensating the aberrations associated with imaging across a tilted coverslip. Additionally, while one could compensate for aberrations in both the imaging and excitation objectives we choose to focus only on the aberrations in the imaging path and therefore use a lower NA air objective for excitation. (b) The geometry of the imaging system. The ray of incidence (green line) crosses the upper surface of a coverslip; the normal vector 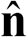 of the coverslip plane is tilted by angle *γ* from *z*-axis in the *x*-*z* plane. The ray is refracted at the interface, travels inside the coverslip(red line), exits from the bottom interface and propagates in the original direction. Without the coverslip, the ray is supposed to propagate along the dashed line.

Fortunately, adaptive optics (AOs), such as deformable mirrors [10] and spatial light modulators [11], have provided a promising approach for complex aberration correction in biomedical imaging [12]. An AO device can add a prescribed phase distortion to actively compensate a distorted wavefront. In biological imaging, it is widely recognized that thick specimens can cause substantial aberrations due to spatial variations of refractive index. In the past two decades, AO devices have been implemented to address this problem of aberration in a variety of microscopy methods, including confocal microscopy [13, 14], wide-field microscopy [15], two-photon microscopy [16, 17], single-molecule imaging [18, 19] and SPIM [20, 21]. However, most existing works have focused on specimen-induced aberrations; system-induced aberrations, which can reflect alignment imperfections or non-ideal sample mounting methods, remains a less discussed aspect with the notable exception of Ref. [20] which addresses the aberrations when imaging across a tube holding the sample. We note that some existing SPIM systems that also have an inverted microscope geometry do not suffer from the same system-induced aberrations as our open-top SPIM setup [7,22,23]. This is due to those other techniques using a single objective beneath the sample that is oriented normal to the coverglass whereas our objectives are oriented at an approximately 45° angle to the coverslip.

Here, we extend the open-top SPIM concept to the case of a high-NA imaging objective with the assistance of AO techniques. Since the majority of the overall aberration is inherent in the system, we carry out a theoretical analysis to estimate its magnitude, spatial structure and major components. Then, using a phase-retrieval method, we experimentally characterize the optical aberration in the system and use a deformable mirror for correction. To test the imaging capability of the system on biological samples, we imaged fluorescently labeled cells with the preconfigured counter-aberration pattern added on the deformable mirror. Even without sample-specific corrections, considerable improvement on the resolution was achieved.

## 2. An estimation of coverslip-induced optical aberration

### 2.1. Theoretical analysis

In this section we evaluate optical aberrations induced by the presence of a tilted coverslip between a point source and a water-dipping objective. We quantify this aberration by calculating the phase shift resulting from the tilted coverslip. The result is used in section 2.2 to determine the nature of the aberration and if it can be addressed using adaptive optics. We assume that the coverslip has #1.5 (170 *μ*m) thickness, which is the case in our imaging experiments. Figure 1(b) illustrates the geometry of the imaging system, wherein the optical axis of the objective is aligned with the *z*-direction. We assume that the point light source is located at the center of the field of view (FOV) and its emission is isotropic. The phase change on the pupil plane caused by a displacement of the point source from the center of FOV can be represented by the first 4 Zernike modes, and thus does not introduce additional optical aberrations.

The maximum cone of light that can enter the imaging system is defined by the numerical aperture (NA) of the objective:

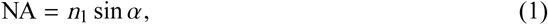

where *n*_1_ is the refractive index of the immersion medium. Assume that the coverslip has thickness *d*, refractive index *n*_2_, and orientation 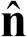. Without loss of generality, we can let 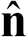 be contained in the *x*-*z* plane, i.e.,

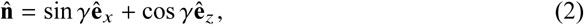

where *γ* is the tilt angle of the coverslip with respect to the *x*-*y* plane. Within the collection cone of the objective, the direction of a light ray, 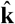, can be specified by the azimuth angle *θ* and the polar angle *φ*:

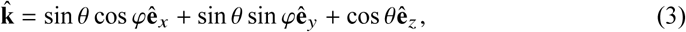

and *θ* satisfies the condition *α + π*/2 ≤ *θ* ≤ *π*. On the upper water-glass interface, the angle of incidence, *ϕ*, is determined by

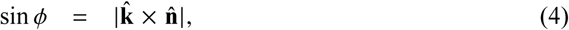

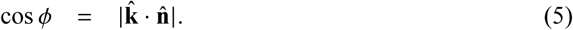

In addition, the *s*- and *p*- directions with respect to the plane of incidence is determined by

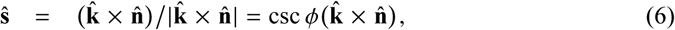

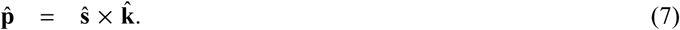

According to Snell’s law, the angle of refraction, *ϕ′*, is associated with *ϕ* by

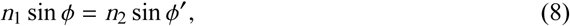

and *ϕ′* can be calculated by substituting Eq. (5) into Eq. (8). Let *β* = *ϕ* − *ϕ′*, then in the 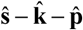 coordinates, the direction vector of the refracted ray, 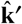 can be denoted by

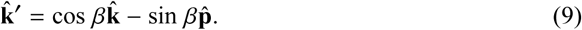

The upper water-coverslip interface can be described by the equation of plane

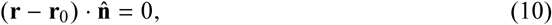

where **r**_0_ is an arbitrary point in the plane. If we set the origin at the point of incidence, then **r**_0_can be set as **0**, and Eq. (10) is simplified as

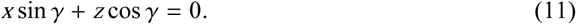

The lower coverslip-water interface, which is translated from the upper interface in the direction of 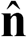 by −*d*, can be described by the equation of plane

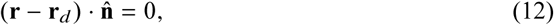

where 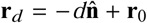. Substituting Eq. (11) into Eq. (12), the latter can be simplified as

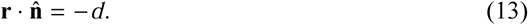

Furthermore, the refracted ray can be described by the equations of a line

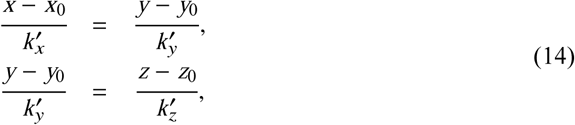

where **r**_0_ is the point of incidence, and 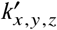 are the *x*, *y*, *z* components of 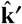 obtained in Eq. (9). Combining Eq. (13) and Eq. (14), the intersection between the lower interface and the refracted ray, i.e., the exit site of the refracted ray, **r**_*e*_ = [*x*_*e*_, *y*_*e*_, *z*_*e*_]^*T*^, can be determined by solving the linear system

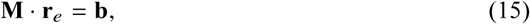

where

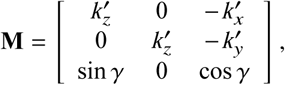

and **b** = [0, 0, −*d*]^*T*^. The optical path length (OPL) of the refracted ray inside the coverslip is

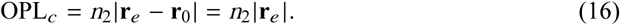

As depicted by Fig. 1, below *z*_*e*_, i.e., the height of the intersection **r**_*e*_, the refracted ray and the virtual original ray (the dashed line) will travel in parallel over the same distance in the immersion medium before they reach the front lens of the objective. This equal distance does not introduce extra aberration, however, the lateral displacement of the ray is equivalent to a displacement of the point source on the sample plane, and thus increases the optical path by

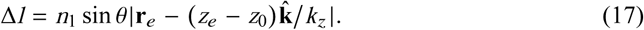

Adding the OPL contributions in Eq. (16) and Eq. (17), the total OPL of the refracted ray is

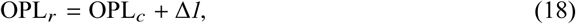

while from *z* = 0 to *z* = *z*_*e*_, the aberration-free OPL should be

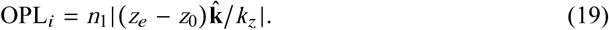

Therefore, the overall phase shift measured by the wavelength is

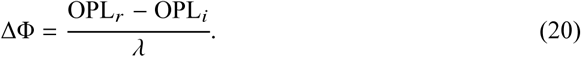

### 2.2. Simulation

Based on the derivations in 2.1, we simulate optical aberrations on the pupil plane with different coverslip tilt angles. The NA of the objective is set at 0.8 or 1.0, and *n*_1_, *n*_2_ are set at 1.33, 1.52, respectively. Figure 2 shows the total, or absolute, aberrations at the pupil plane as well as the aberrations at the pupil plane relative to the untilted coverslip at 0°, 15°, 35° (maximum practical tilt for a 1.0 NA objective because of geometric constraints), and 45° (maximum practical tilt for a 0.8 NA objective). In each case, the phase shift as a function of *θ* and *φ* is calculated using Eq. (20); the tilt and piston components are extracted from the x cross section and subtracted from the raw pupil function. In addition to the coverslip-induced spherical aberrations already present at 0° tilt, calculation results show an increasing amount of phase aberrations with higher tilt angles. At both 35° (for 1.0 NA objective) and 45° (for 0.8 NA objective) coverslip tilt, the steepest change in phase aberrations occurs at the rightmost edge of the pupil plane, where light rays hit the coverslip at the highest incident angles. Other than this edge, the phase aberrations are rather smooth and well suited for correction with a deformable mirror (DM).

**Fig. 2.**
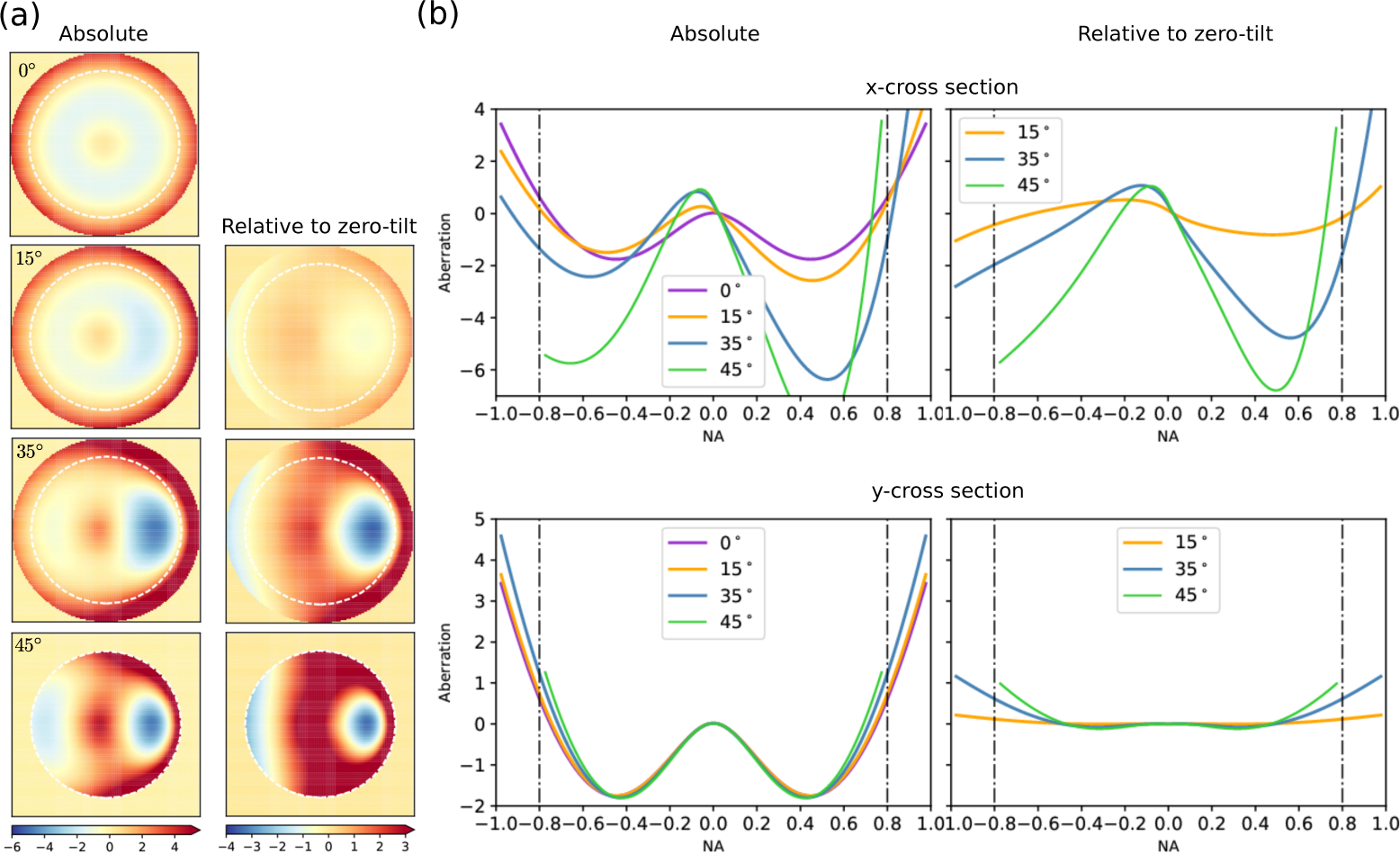
(a) Optical aberrations on the pupil plane of a NA=1.0 objective under 0°, 15°, 35°, 45° coverslip tilt. The linear non-aberrational components (piston, tip/tilt) are subtracted from the phase difference calculated with Eq. (20). The white dashed circle represents 0.8 NA. At 45° coverslip tilt, the pupil function is cropped to NA = 0.8 because of objective geometric constraint. The left column shows the absolute aberration in each case, and the right column shows the relative aberration with respect to that in the zero-tilt case in order to remove the spherical aberrations resulting from imaging across a glass coverslip. Color bar unit: wavelengths. (b) Cross sections of the absolute (left column) and relative (right column) aberrations. Unit: wavelength.

Besides phase aberrations discussed above, light intensity is diminished due to reflections at the water-coverslip interface. The power loss is negligible at small angles of incidence, but can be significant at large angles of incidence; the latter would occur for a large coverslip tilt angle. At each interface, the reflectances for *s*-polarized light and *p*-polarized light, *R*_*s*_ and *R*_*p*_, can be determined by the Fresnel equations:

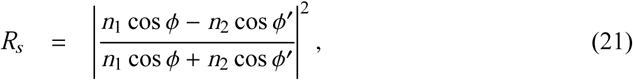

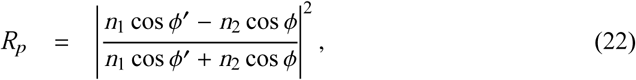

where *ϕ*, *ϕ′* are the angles of incidence and refraction, respectively. Suppose that the incident light is unpolarized, i.e., *P*_*s*_ =*P*_*p*_ =*P*_0_/2, then the transmitted power after crossing the two interfaces is

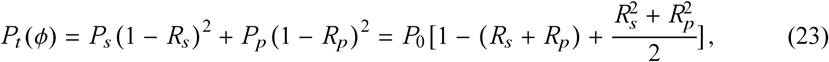

so the total transmittance through the coverslip is

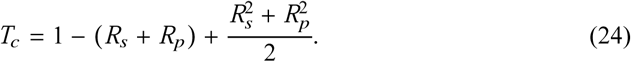

Figure 3 shows *T*_*c*_(*θ*, *φ*) on the pupil plane of an NA = 0.8/1.0 objective for the same four tilt angles in Fig. 2. It can be seen that while the transmittance within most of the pupil plane was close to 1 in all cases, at 35° (for 1.0 NA objective) and 45° (for 0.8 NA objective) coverslip tilt, the amplitude loss on the rightmost edge of the pupil plane is significant. This effect caused an effectively small clipping of the pupil plane and hence a small loss of NA. On the other hand, this reduction of transmittance exactly coincides with the part of pupil plane containing the sharpest variation of phase aberrations. Consequently, imperfect phase aberration correction at this rightmost edge of the pupil plane should not have a large effect on the imaging performance.

**Fig. 3.**
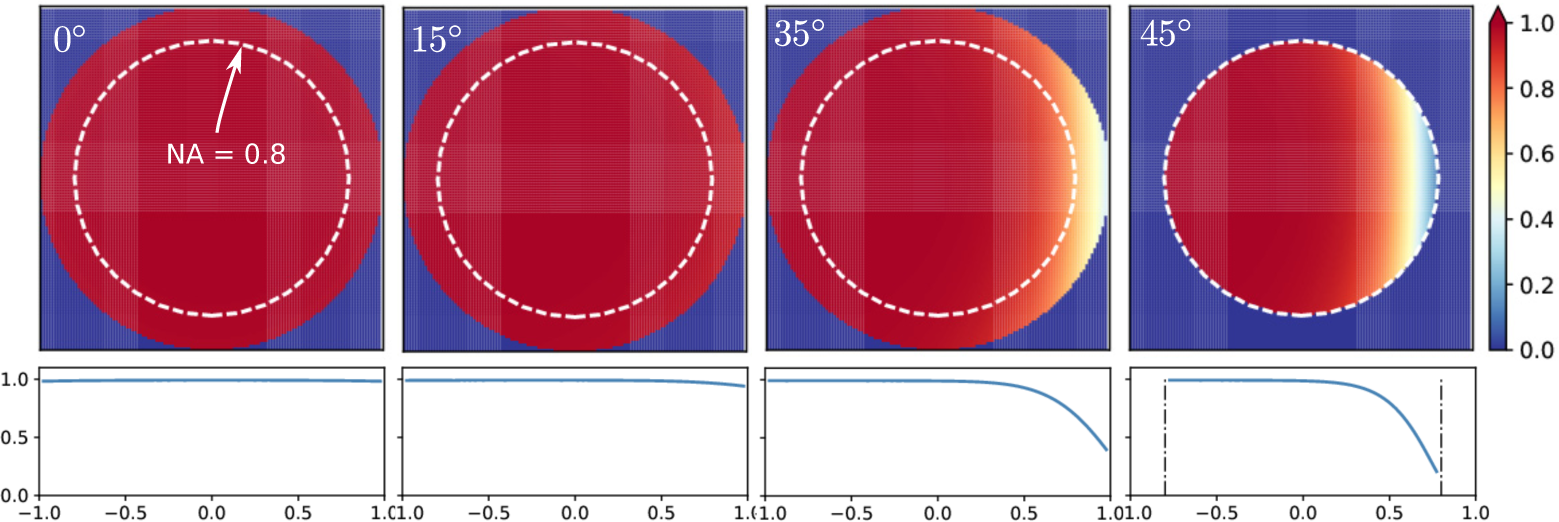
Total transmittance of rays within the collection cone at *γ* = 0°, 15°, 35°, 45° degrees, calculated for NA = 0.8 (white, dashed circle) and 1.0 (full circle). The cross sections along the x-direction are shown in the bottom row.

## 3. Experiment

### 3.1. Construction of the AO-SPIM

To test the high-NA objective performance when imaging across a tilted coverslip, we layout our SPIM system horizontally on an optical table in order to more easily test different high-NA objectives and tilt angles of the sample (Fig. 4). A long working-distance 20× 0.42 NA air objective (Mitutoyo) in combination with an *f* = 200 mm cylindrical lens (Thorlabs) are used for excitation; a 40× 0.8 NA infinity-corrected, water-dipping objective (MRD07420, Nikon), is used for imaging. With this objective combination, the 0.8 NA imaging objective can operate at a coverslip tilt angle of 40°. The collimated laser beam forms a light-sheet after passing the excitation optics. By co-focusing the two objectives, the light-sheet can be confined to the focal plane of the high-NA objective.

**Fig. 4.**
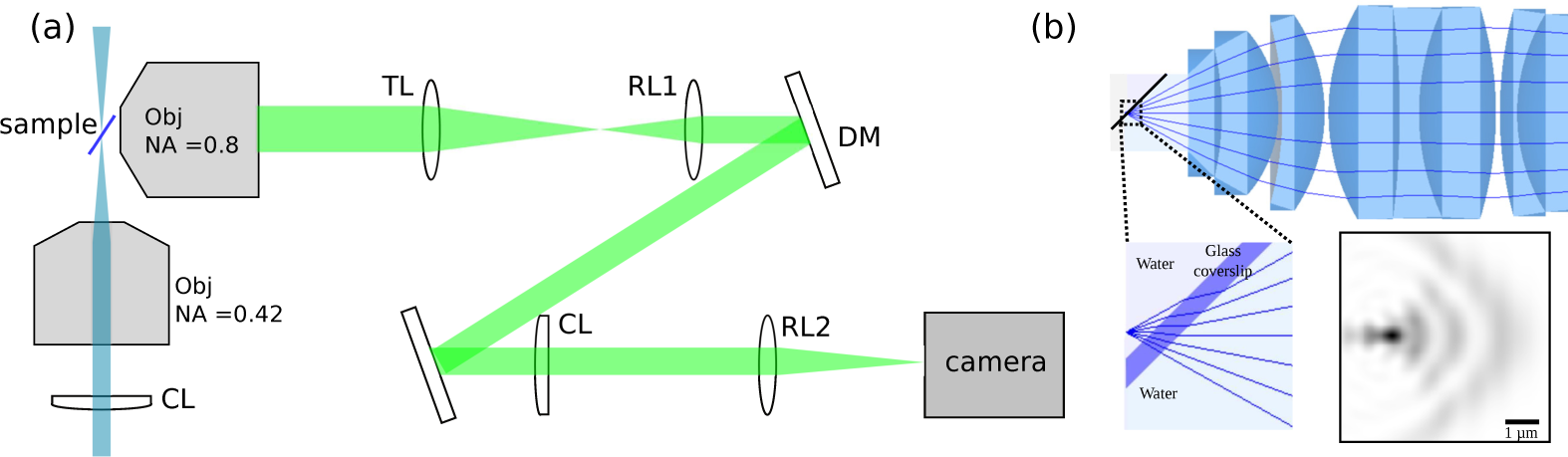
(a) Experimental setup. (b) Results of simulating our optical design with Zemax. We simulate a point-source situated behind a coverglass tilted around 40° to the optical axis. The first few optical elements of a 40× water dipping objective are shown. Light rays are shown emerging from the point source and being refracted at the water-glass and glass-water interfaces. The simulated point spread function is shown. We incorporated a cylindrical lens downstream from the objective in this simulation to counter some of the aberrations.

To accommodate the water-dipping objective, we constructed a small water-containing cube (dimensions 3.5 cm × 3.5 cm × 3.5 cm). The imaging objective dips into the chamber from one side and is sealed with an O-ring. The excitation light enters through a glass window on a perpendicular side. The top of the chamber is open to allow the sample to be inserted from above. The sample is either attached to a coverslip or held between a glass slide and coverslip and held perpendicular to the optical table. By using a manual rotation stage we can tilt the slide or coverslip at a known angle with respect to the optical axis. A motorized translation stage (MT1-Z8, Thorlabs) allows us to move the sample along the optical axis of the imaging objective. The excitation light is provided by a 473 nm diode-pumped solid state laser (MBLFN-473-50mW, Changchun New Industries Optoelectronics, China). Images are magnified by a pair of relay lenses and captured by a scientific CMOS camera (Flash 4.0, Hamamatsu). In all cases, we used #1.5 (170 *μ*m) coverslips.

In our previous low NA open-top SPIM system, the system-induced optical aberration is partially corrected by a 10 m focal length cylindrical lens (SCX-50.8-5000.0, Melles-Griot) placed before the second relay lens. A Zemax simulation of the point-spread function (PSF) at the focus shows that the optical aberration using the 0.8 NA objective, even in the presence of the counter-aberration cylindrical lens, is severe when the tilt angle is 40°. To correct the rest of the aberration, we place a 140-actuator, continuous surface deformable mirror (Multi-DM, Boston Micromachines) at the conjugate plane of the objective back pupil. Each actuator has a maximum stroke of 3.5 *μ*m, which is equivalent to 7 wavelengths for fluorescent signals around 500 nm. The conjugate pupil diameter is reduced to 4 mm by the telescope formed by the tube lens (TL, *f* = 200 mm) and the first relay lens (RL1, *f* = 100 mm) to match the active area (4.4 mm diameter) of the deformable mirror.

### 3.2. Aberration measurement and correction

In order to experimentally characterize the aberration in the present setup, we record the PSFs of sub-diffraction limit sized fluorescent beads to numerically retrieve the pupil function [24]. We use 200-nm diameter fluorescent beads that are either attached to a glass coverslip or embedded in 1% agarose gel sandwiched between a coverslip and a glass slide. The sample is then stepped in the z-direction as depicted in Fig. 4. Unless otherwise noted, the PSFs are acquired in the epi-illumination mode where the excitation and emitted light both pass through the same objective. This allows us to capture a full three dimensional PSF unbiased by the intensity distribution of the light sheet in the z direction. Typically, we record 21 images separated by Δ*z* = 0.5 *μ*m of a single bead to find the pupil function using the iterative phase-retrieval method previously described [24, 25].

An example of a pupil function retrieved for a coverslip tilt angle around 40° is shown in Fig. 5(a). The phase retrieval procedure is iterated several times (seven times in the case shown in Fig. 5(a)) to produce a final pupil function. Each iteration consists of measuring the three-dimensional PSF of a fluorescent bead, retrieving the pupil function and updating the deformable mirror. These multiple iterations are needed as a single phase-retrieval process does not always, especially in the presence of large aberrations with multiple phase wraps, produce error-free pupil functions. Additionally, the multiple iterations mitigate the effects of uncertainty in the relationship between the voltage applied to the deformable mirror’s actuators and their displacement and the effects of inter-actuator coupling. The retrieved pupil function is reversed and discretized into 12×12 segments as shown in Fig. 5(b), then scaled to a voltage to be applied to the deformable mirror for aberration correction. The associated Zernike coefficients of modes 5–22 are shown in Fig. 5(d). At the end of the 7 iterations, the overall range of voltages applied on the deformable mirror segments takes 69% of the mirror stroke, which is equivalent to 4.7 wavelengths. The retrieved pupil function has little aberration except on the rightmost edge (Fig. 5(c)), and the RMS error over the pupil plane is 0.21 wavelength.

**Fig. 5.**
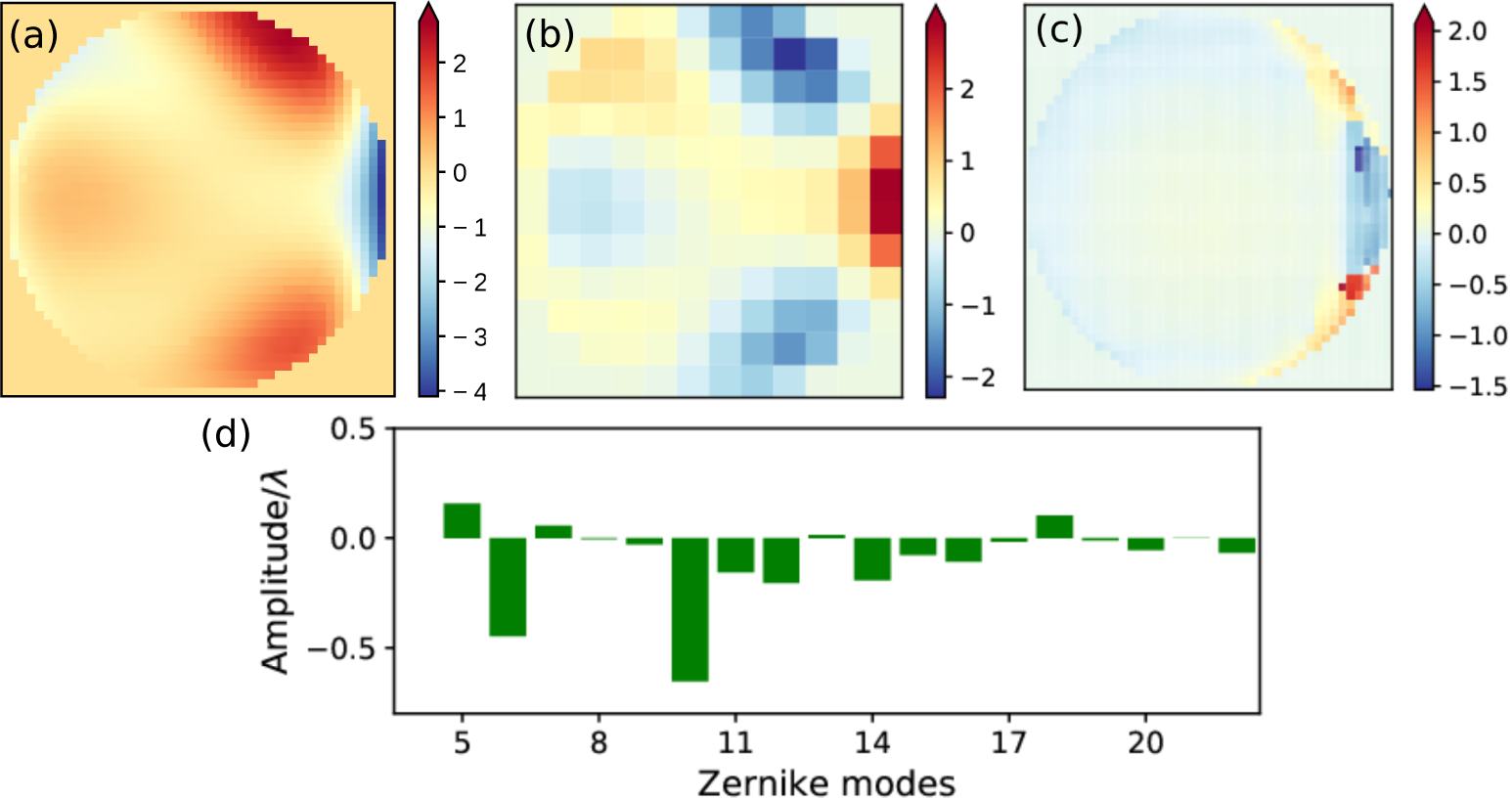
(a) The final pupil function retrieved after 7 iterations. In each iteration, the retrieved pupil function is reversed and discretized into 12×12 segments, then scaled up and applied to the 140-segment deformable mirror. The overall correction pattern added on the deformable mirror is shown in (b). (c) With the correction in (b) applied to the deformable mirror, a majority of the system aberration is eliminated except at the rightmost edge of the pupil plane as shown in this retrieved pupil function. (d) The coefficients of 5–22 zernike components in the retrieved system aberration (a).

Figures 6(a)-6(c) shows orthogonal slices through the three-dimensional point-spread function when imaging across an approximately 40°-tilted coverglass. The cylindrical lens is in place in the detection path to initially correct as much of the aberrations as possible. Fitting a Gaussian function to a line profile though this PSF yields a full-width at half maximum (FWHM) of 4.6 *μ*m along the long axis of the PSF, 0.5 *μ*m in x, and 0.6 *μ*m in y directions (see Fig. 6(j)). After six iterations of phase-retrieval and correction with the deformable mirror the PSF quality is significantly improved, as shown in Figs. 6(d)-6(f). With this correction, the FWHM of the Gaussian-fitted PSF is 2.6 *μ*m in z and 0.4 *μ*m in both x and y (Fig. 6(k)). The x-y PSF width is close to the theoretical ideal for a 0.8 NA objective, indicating a near-diffraction-limited performance. We noticed a slight tilt of the PSF in the xz cross section, possibly resulted from the clipping of the pupil plane at its rightmost edge due to reduced transmission. In addition to the proof-of-principle setup using epi-illumination, we also measured the PSF using light-sheet illumination, i.e., illuminating the sample via the path depicted in Fig. 4 that is orthogonal to the imaging objective. As shown in Figs. 6(g)-6(i), due to the reduced illumination depth of the light sheet, the axial extent of the PSF is further reduced. The axial FWHM of the Gaussian-fitted PSF with the light-sheet illumination is 1.9 *μ*m (figure not shown). In addition to measuring the shape of the PSF before and after correction we have also measured how the signal from fluorescent beads changes. After correcting for system-induced aberrations we measured the intensity from 25 fluorescent beads none of which were the bead used in determining the correction to apply to our deformable mirror. The average factor by which the signal increased upon correcting the aberrations was 2.6 with a standard deviation of 0.2.

**Fig. 6.**
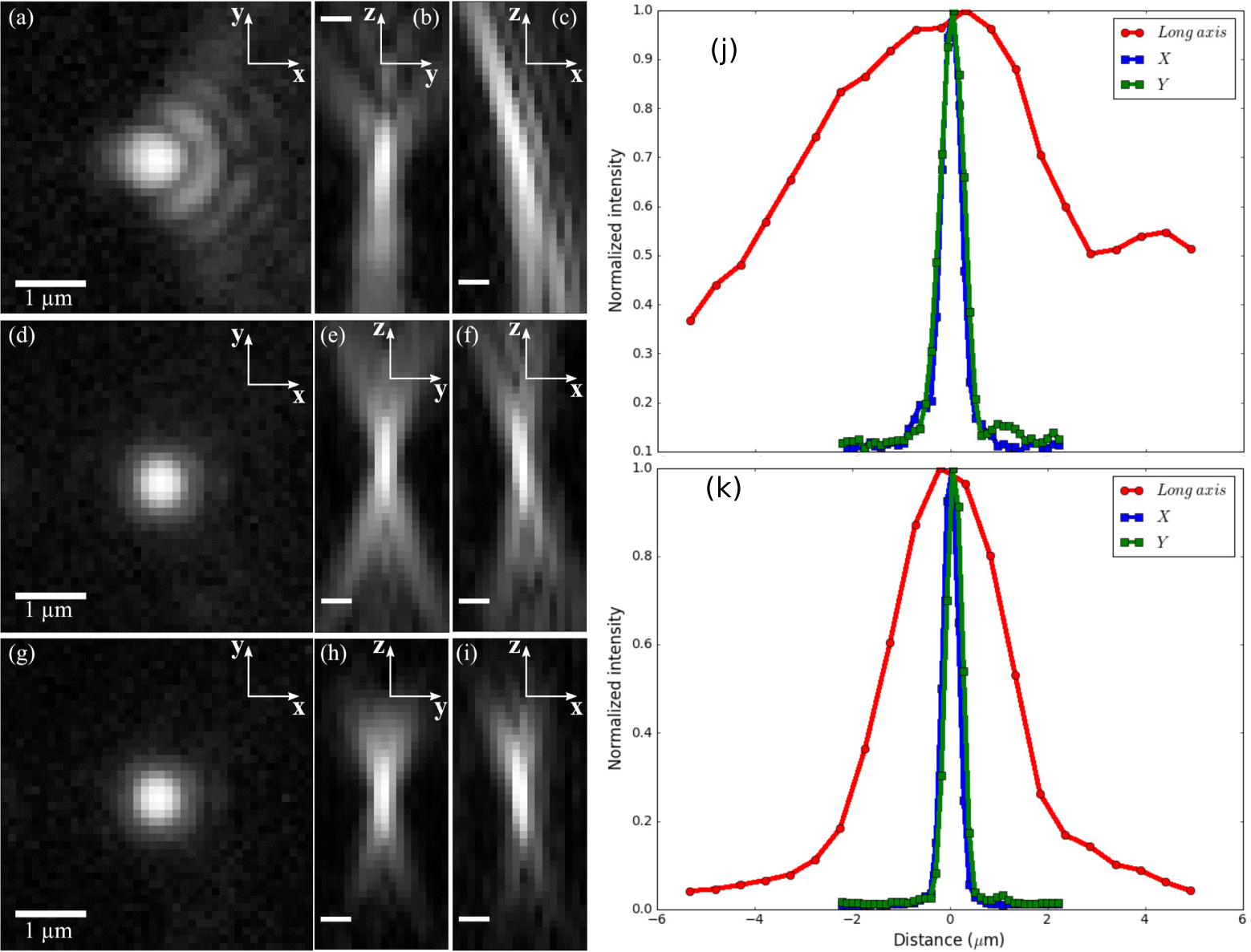
The PSF of our microscope without and with aberration corrections from the deformable mirror, all displayed with log scaling. All scale bars are 1 *μ*m. In (a)-(c), only the cylindrical lens placed in the imaging path corrects some of the aberrations. The deformable mirror is flat. In (d)-(f), a pattern is placed on the deformable mirror found through iteratively finding the pupil function and adjusting the deformable mirror. In (g)-(i), a sheet of excitation light coming from an orthogonally placed objective is used rather than using wide-field illumination as in the previous cases. (a), (d), and (g) display slices of the PSF in the x-y plane. (b), (e), and (h) display slices in the z-y plane. (c), (f), and (i) display slices in the z-x plane. (j), (k): Cross sections of the PSFs in x, y, and z directions before and after the aberration correction.

Although the air-water interface in the excitation path introduces aberrations into the excitation path and degrades the quality of the light sheet, we have not yet attempted to correct these aberrations. As the excitation light sheet is formed with a much lower NA objective (0.42 NA in our case) the aberrations will be much less severe. However, additional efforts on the excitation side would be necessary to achieve a diffraction-limited thickness of the light sheet.

We compared PSFs measured for fluorescent bead samples prepared in the two approaches, i.e., beads are directly attached to the backside of a coverslip and embedded in agarose gel, respectively. Over a volume extending several tens of microns in all dimensions, we do not observe any significant difference between the PSFs in the two cases. However, when one images farther from the point where the PSF is acquired to retrieve the patterns added on the deformable mirror, we expect the performance of the microscope to degrade. We plan to determine the size of the isoplanatic patch in the future.

### 3.3. Imaging biological samples

To demonstrate the biological applications of our high-NA open-top SPIM, we first imaged human embryonic kidney 293 (HEK) cells with fluorescently labeled nuclear lamina with the setup depicted in Fig. 4(a). Like the fluorescent-bead samples, the cells were placed between a glass slide and a coverslip, with the latter facing and at approximately 45° to the imaging objective (40× 0.8 NA). The excitation light sheet was provided by an air objective (20× 0.42 NA). Image stacks were acquired by moving the sample in 1 *μ*m steps along the optical axis of the imaging objective. Immediately prior to imaging the cells, the pupil function under the specific tilt angle of the sample had been characterized with 200 nm fluorescent beads, and the deformable mirror was adjusted accordingly. Figure 7 shows images of a few cells. With the correction applied, the image exhibits sub-micron resolution even though the correction only compensates system-induced aberration while leaving the sample-induced aberrations unaddressed. Structures such as nuclei and cell membranes can be reliably resolved at this stage.

**Fig. 7.**
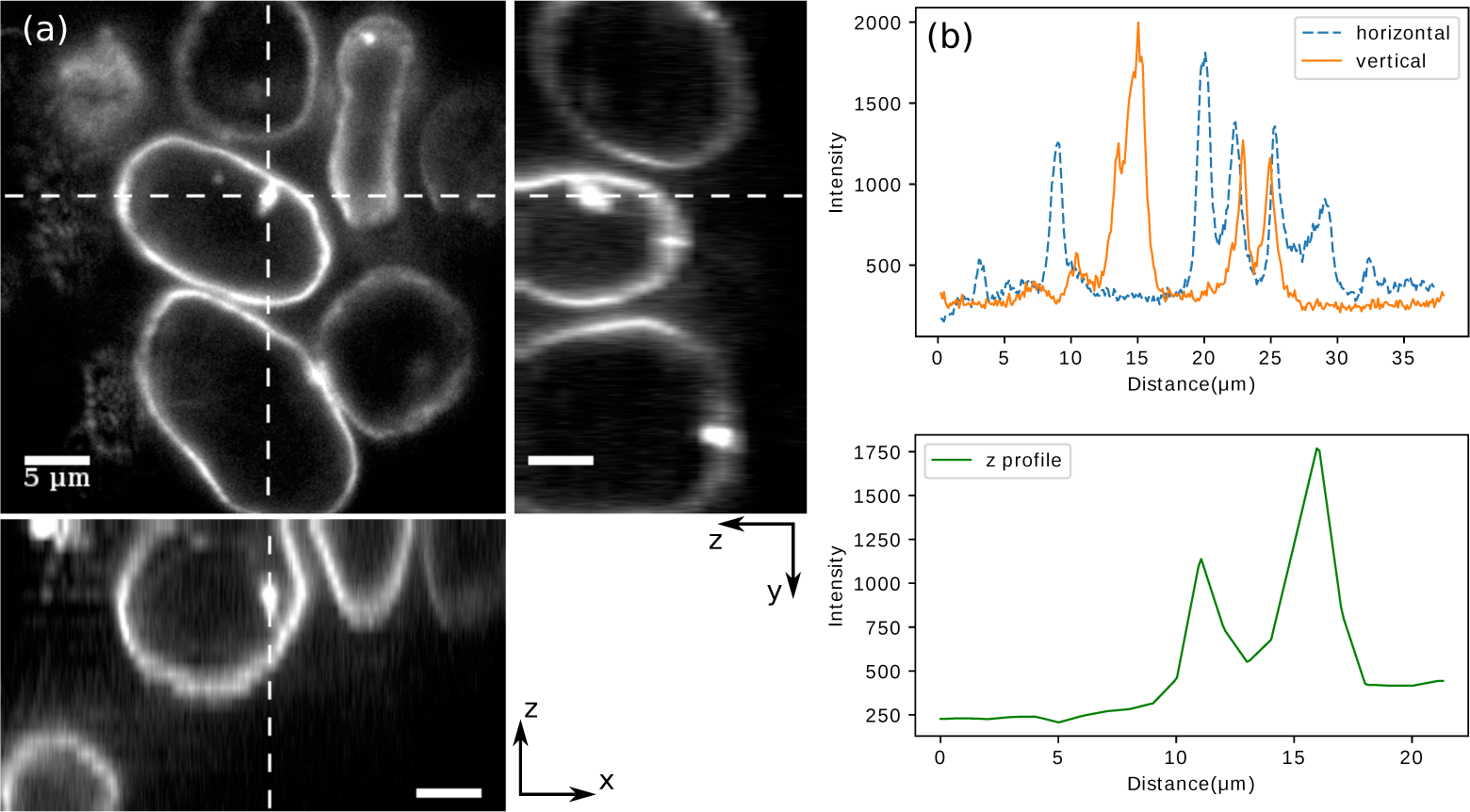
HEK cells with fluorescently labeled nuclear lamina were imaged with our SPIM using a cylindrical lens and a deformable mirror to reduce aberrations. (a) Upper left: an xy frame of the acquired stack. Upper right and bottom: y-z and x-z view of the slices along the dashed lines. The displayed volume covers 22 *μ*m along the z-axis. (b) Upper panel: the intensity profiles along the horizontal and vertical dashed lines, respectively. Lower panel: the z-profile along the cross section.

Next, we tested our methods on a horseradish peroxide (HRP)-stained *Drosophila* embryo. Image stacks were captured using the same acquisition strategy as it was for HEK cells; however, instead of calculating the deformable mirror pattern afresh via the phase-retrieval routine, we directly applied to the mirror a pattern configured previously under the same tilt angle. As a comparison, an image stack across the same sample region was acquired with the deformable mirror “flattened”. Figure 8 shows a sample frame with and without the mirror correcting system-induced aberrations. As can be seen in both the images and profiles of the selected line cuts, a significant improvement of resolution was brought about by the prescribed correction.

**Fig. 8.**
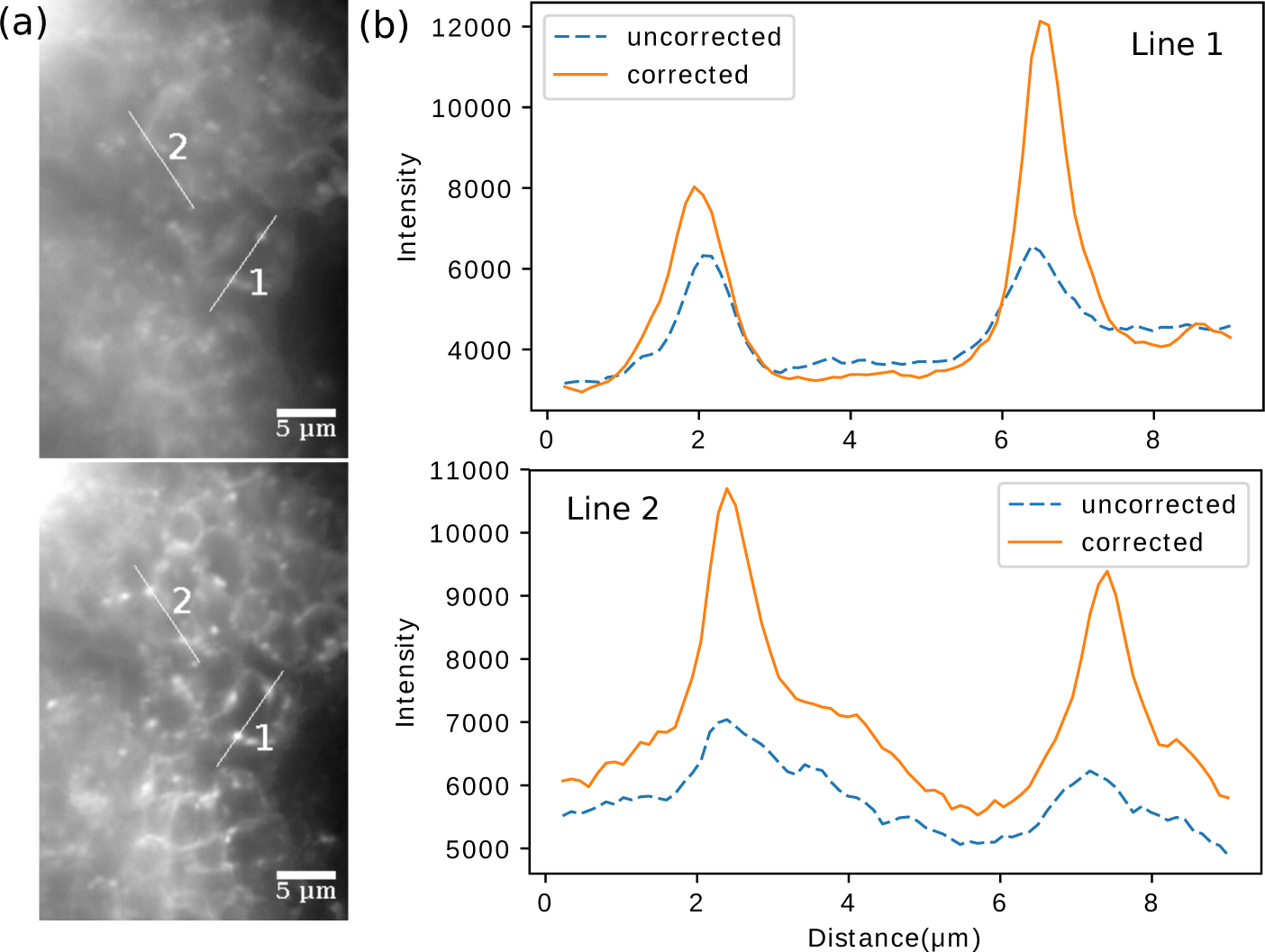
Images of a *Drosophila* embryo before and after a preconfigured correction pattern was applied on the deformable mirror. Two 8-*μ*m line cuts (1 and 2) are made across the same structures in both cases, and their intensity profiles are shown in (b).

We noticed that switching out the fluorescent bead sample used for aberration characterization for the sample of labeled cells invariably alters, however slightly, the tilt angle of the coverglass, the coverglass thickness and the distance of the sample beyond the coverglass. However, we found that the slight variations in these parameters do not significantly change the necessary pattern on the deformable mirror to obtain high-resolution images. This robustness suggests that the major component of the overall aberration is system-induced and remains static given the alignment is preserved. Thus, for samples that do not vary largely in composition over the whole volume, e.g., single layers of fixed cells or cleared tissues, the correction pattern calculated from a point source can be directly applied to the deformable mirror or, alternatively, a custom static optical element for system-induced aberration correction can be employed. This “once and forever” mode would no longer suffice for samples that change drastically over time and space, e.g., developing fly embryos [26]. Faster computational routines would be required to dynamically shape the correction pattern, and multiple AO devices may be needed in the optical path for complementary optical corrections [2].

## 4. Conclusion

We have demonstrated the implementation of a high-resolution open-top SPIM system, which allows one to use specimen prepared in conventional sample holders. Central to the new system is a high-NA objective that collects signals through a tilted coverslip. Although this configuration introduces considerable aberration into the images, we have shown that such aberration can be reliably retrieved by measuring emissions of a point source, and then be largely reversed by adding an opposite aberration pattern on a deformable mirror. Simulations of the imaging system and experimental results both suggest that uncorrected aberrations introduced by the tilted coverslip between the objective and the sample primarily exists at one edge of the exit pupil. This design provides an optically simple and versatile approach to image high-content samples at sub-micron resolution on a SPIM platform.

## Acknowledgments

The authors would like to thank Dr. Sayaka Sekine for kindly providing the HEK cells, and Dr. Daichi Kamiyama for providing the *Drosophila* embryo.

